# Gamma Entrainment Improves Synchronization Deficits in Dementia Patients

**DOI:** 10.1101/2021.09.30.462389

**Authors:** Mojtaba Lahijanian, Hamid Aghajan, Zahra Vahabi, Arshia Afzal

## Abstract

Non-invasive gamma entrainment has shown promising results in alleviating cognitive symptoms of Alzheimer’s disease in mice and humans. In this study, we examine improvements in the synchronization characteristics of the brain’s oscillations induced by 40Hz auditory stimulation based on electroencephalography data recorded from a group of dementia patients. We observed that when the quality of entrainment surpasses a certain level, several indicators of brain synchronization significantly improve. Specifically, the entrained oscillatory activity maintains temporal phase stability in the frontal, parietal, and occipital regions, and persistent spatial phase coupling between them. In addition, notable theta-gamma phase-amplitude coupling is observed in these areas. Interestingly, a high theta power at rest predicts the quality of entrainment. We identify differentiating attributes of temporal/spatial synchronization and cross-frequency coupling in the data of two groups with entrained and non-entrained responses which point to enhanced network synchronization caused by entrainment and can explain its potential therapeutic effects.

## Introduction

As a neurodegenerative disease, dementia due to Alzheimer’s disease (AD) involves the brain via different mechanisms. The accumulation of insoluble plaque called amyloid-beta (Aβ) outside neurons, and neurofibrillary tangles called phosphorylated tau inside neurons, are known as AD’s major characteristics which lead the brain to a more deteriorative state and cause loss of synapses^1–3^. As these neuropathological effects advance, signs of network operation deficiencies appear. Brain rhythms get out of order, and neuronal synchrony of brain oscillations becomes disheveled, causing structural and functional changes in the brain’s neuronal networks^4^. As a major manifest of network disruption, the efficacy of inhibitory gamma-band activity diminishes^5–8^, excitatory neurons act superfluously^1,9,10^, leading to further acceleration of Aβ production as a byproduct of neural activity^11^ and further advancing the cycle of the disease progress. The quality of cross-frequency coupling functions, such as the theta-gamma coupling which plays a key role in cognitive tasks and memory retrieval operations^4,6,12,13^, is also known to be affected as network disruptions continue^14,15^. Much of oscillatory abnormalities associated with AD commence in areas such as the entorhinal cortex, the limbic system, and the hippocampus, which are responsible for cognition, memory tasks, and sensory processes, and known to involve high-frequency oscillatory activity in the gamma band^6,9,16^.

Neurodegeneration, network alteration, and cognitive deficit are all manifested in AD^4,5,15^, which informs the potential correlation that exists between them. Although definite cause and effect links governing the relationships among these phenomena need to be yet clarified, much of the initial approaches to AD treatment have targeted neurodegeneration or symptom relief through pharmaceutical remedies. Most notably, the newly FDA-approved Aducanumab^17^ is reported to reduce the Aβ load in the brain. However, its much debated clinical trials have fallen short in proving noticeable benefit in improving cognitive symptoms of dementia^1,18^, and even its efficacy in lowering Aβ is questioned^19^, extending the controversy about it to a span from potential safety risks and high cost to the accelerated approval process that relied on this surrogate endpoint and not cognitive improvements^20,21^. It has also been reported earlier that while reducing the Aβ load can slow the progression of AD, even a great reduction in its level may not lead to memory retrieval improvement^1^. The shortcomings of the current remedial approaches to AD treatment may stem from ignoring the intricate interactions between the local synaptic damage inflicted by Aβ aggregation and the large-scale network alterations that shift the operating point of feedforward and feedback neuronal networks.

It is still a matter of debate whether network alterations such as deficits in gamma activity are a consequence of the underlying neurodegenerative processes or they indeed play a causal role in inducing neuropathological changes that promote the disease progression. While neuropathological effects such as the aggregation of surplus Aβ in intercellular space and the ensuing synaptic loss are reported to cause hyperexcitability in excitatory neural populations leading to network disruption^1,9^, other evidences point to the presence of network alterations such as reduced gamma oscillations or diminished theta-gamma coupling even before plaque aggregates are ostensibly detectable^22–24^. Additionally, more recent evidence suggests a causal direction from the quality of brain oscillations to the performance of memory and attention functions^25,26^. Based on these observations, approaches to mitigate or compensate for network alterations manifested in the lowered quality of the brain’s oscillatory activity can serve as alternative pathways to AD treatment. Detailed examination of the changes in the brain’s oscillatory attributes such as the quality of synchronization or cross-frequency coupling instilled through such approaches can offer insight into the interwoven cycles of cause and effect involving neuropathological changes and network alterations, paving the way for more integrative approaches to the treatment of the disease.

Interventions based on stimulating neural populations in the gamma band have been recently proposed and are under examination on animal models of AD^22,27–29^ and in limited human studies^30–36^. These approaches aim to reinvigorate the functionality of the brain networks through synchronized pumping of rhythmic energy via one or multiple sensory modalities. Stimulating neuronal networks of the brain by an external stimulus at a specific frequency drives the neurons to undergo oscillatory spiking behavior in that frequency, which is referred to as entrainment^37^. The idea behind such network training approaches is to compensate for the functional weaknesses caused by synaptic loss due to Aβ plaque deposition. These methods primarily focus on combating the network alteration characteristics of AD, and early results have shown promising outcomes in improving cognitive functionality of the subjects. For example, optogenetically-induced 40Hz entrainment of the Parvalbumin (PV) inhibitory interneuron cells in the hippocampus has been reported to reduce the level of amyloid plaque in different mouse models of AD^22,29^. Non-invasive entrainment of gamma oscillations using auditory and visual stimulation has been shown to improve memory performance in AD mouse models and more interestingly, reduce the amyloid load in the auditory cortex, hippocampus, and the medial prefrontal cortex of the mice^22,27^. Exposure to flickering light at 40Hz has been shown to produce a similar effect in the visual cortex^22,28,29^. Remarkably, treating the AD mouse models with repeated entrainment sessions has been reported to result in improved memory and cognitive functions and slowing the neurodegenerative effects of AD^22,27–29^. The induction of gamma oscillations in AD mouse models through sensory stimulation has been shown to also improve neuroimmune biochemical signaling known to promote microglia activity^38^ and enhance their Aβ uptake^22^.

In this study, we aim to provide an interpretation of the therapeutic effects of gamma entrainment by examining improvements in the synchronization characteristics of the brain oscillations induced by gamma stimulation. Based on EEG data recorded during 40Hz auditory stimulation of a group of elderly participants suffering from dementia, a metric is defined for the quality of the entrainment response and the participants are divided into entrained and non-entrained groups. We identify attributes of network synchronization caused by entrainment which significantly differentiate the two groups. These attributes consist of temporal, spatial, and cross-frequency coupling characteristics of the response which have been reported in various studies as major oscillatory deficit manifests of network alteration in AD^4,39,40^. Specifically, temporal phase stability during stimulation as well as the strength of spatial phase coupling of different brain regions and the quality of theta-nested gamma oscillations are examined as indicators of synchronized network activity. Enhancements in these indicators due to entrainment reflect the recovery of network synchronization which can help explain cognitive improvements reported for gamma band entrainment.

## Results

### The occurrence of entrainment

There is no standard definition for an entrained brain. While stimulating the brain with 40Hz auditory or visual stimuli is referred to as entrainment, our results show that such stimulation does not necessarily cause notable entrained oscillatory activity in all brains. Based on earlier studies on entrainment^31,33,37^, a clear response of the brain to the stimulant frequency is expected as a peak in the spectrogram of the recorded response on that frequency. Accordingly, we propose a method to determine the occurrence of entrainment (see Methods). Figs. 1a-b illustrate matrices of entrainment occurrence for all participants across all channels and during all trials.

**Figure 1.**
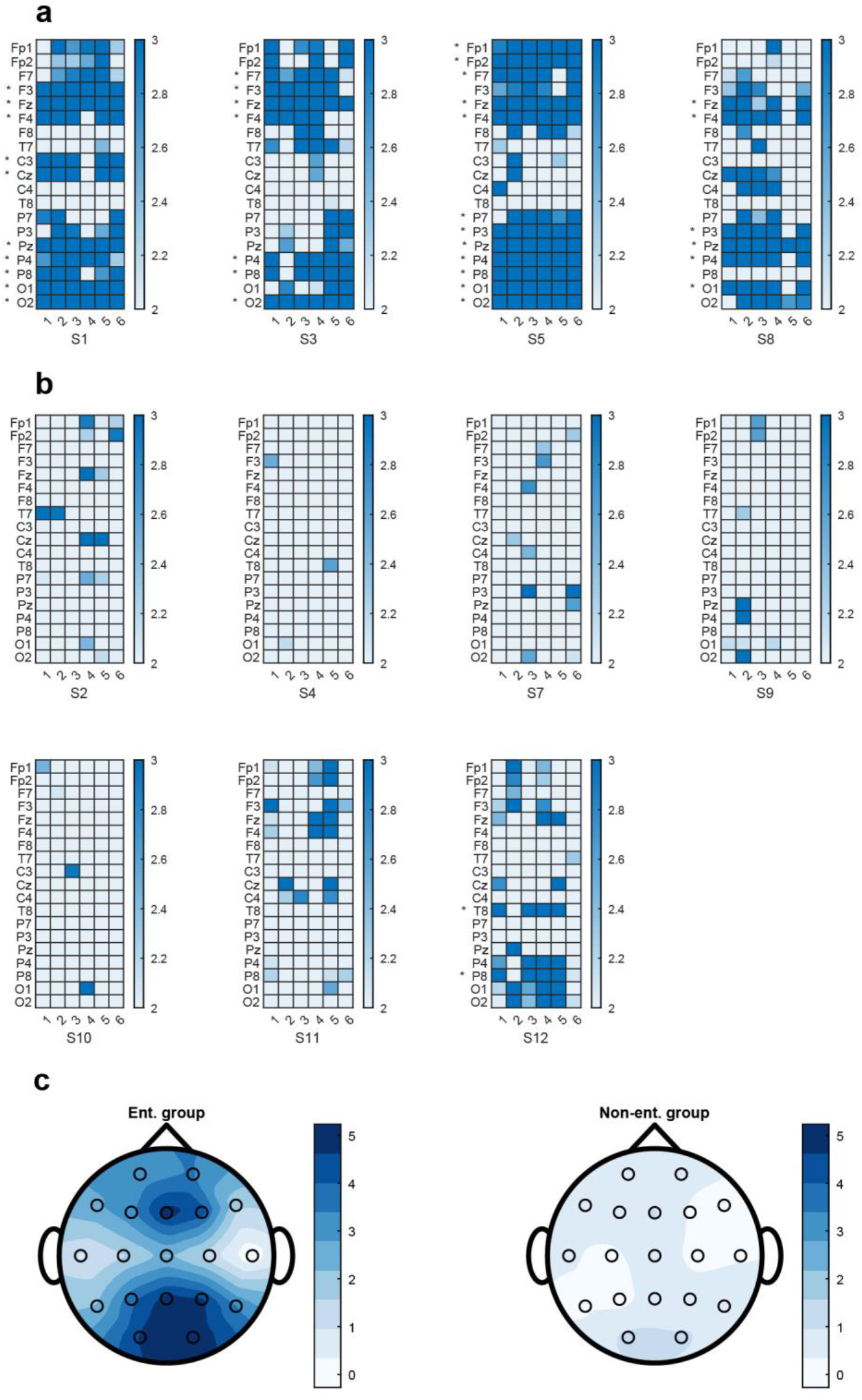
Occurrence of entrainment during stimulation cycles. (**a**) Checkered diagrams show the occurrence of entrainment for all trials (columns), all channels (rows), and all participants in the entrained group (n = 4). The color of each matrix element represents a z-score value of the 40Hz component’s amplitude in the frequency response and dark blue elements indicate entrained responses (z-score above 3). The star next to each channel implies that the majority of trial responses are entrained. (**b**) Same as **a** for the non-entrained group (n = 7). (**c**) Topographic distribution of entrainment over the scalp for the two groups. The gamma entrainment score (trial-averaged z-score, see methods for more information) is color coded and the plots show the average group response to the external stimulus. The frontal, parietal, and occipital channels show highest entrained responses in the entrained group. Ent.: Entrained, Non-ent.: Non-entrained.

We further define a gamma entrainment score to quantitatively assess how well the brain is entrained due to external stimulation. Based on the results shown in Fig. 1c, entrainment is mainly observed in frontal, parietal, and occipital electrodes, with the F3, Fz, F4, P3, Pz, P4, O1, and O2 channels showing highest entrained responses. As the activity of the frontal lobe has been used in entrainment studies as an indication of improved cognitive functions^27–29^, an entrained response at Fz across trials is used to divide the participants into two groups of entrained and non-entrained. Fig. 1c shows the average distribution of entrainment occurrences across channels for the two groups. It is noteworthy that an auditory stimulant alone can drive entrainment in a rather large area of the brain in the entrained group.

### Intrinsic theta power is highly correlated with entrainability of gamma oscillations

To examine the relationship between the power of theta and gamma oscillations on channel Fz, we calculate ensemble averages for the stimulation and rest trials separately. We normalize the power of the target frequency band during stimulation with the power measured during the rest trials.

As Fig. 2a illustrates, the normalized gamma power response on Fz is considerably higher for the entrained group compared to the non-entrained group. This difference is significant (ttest2, t_9_ = 3.0111, p-value = 0.0147; Fig. 2c right) and does also exist between the two groups across most other channels (as evidenced in Fig. 1c). Fig. 2b indicates that the normalized theta power does not significantly differ between the two groups (ttest2, t_9_ = −0.2009, p-value = 0.8452; Fig. 2c left).

**Figure 2.**
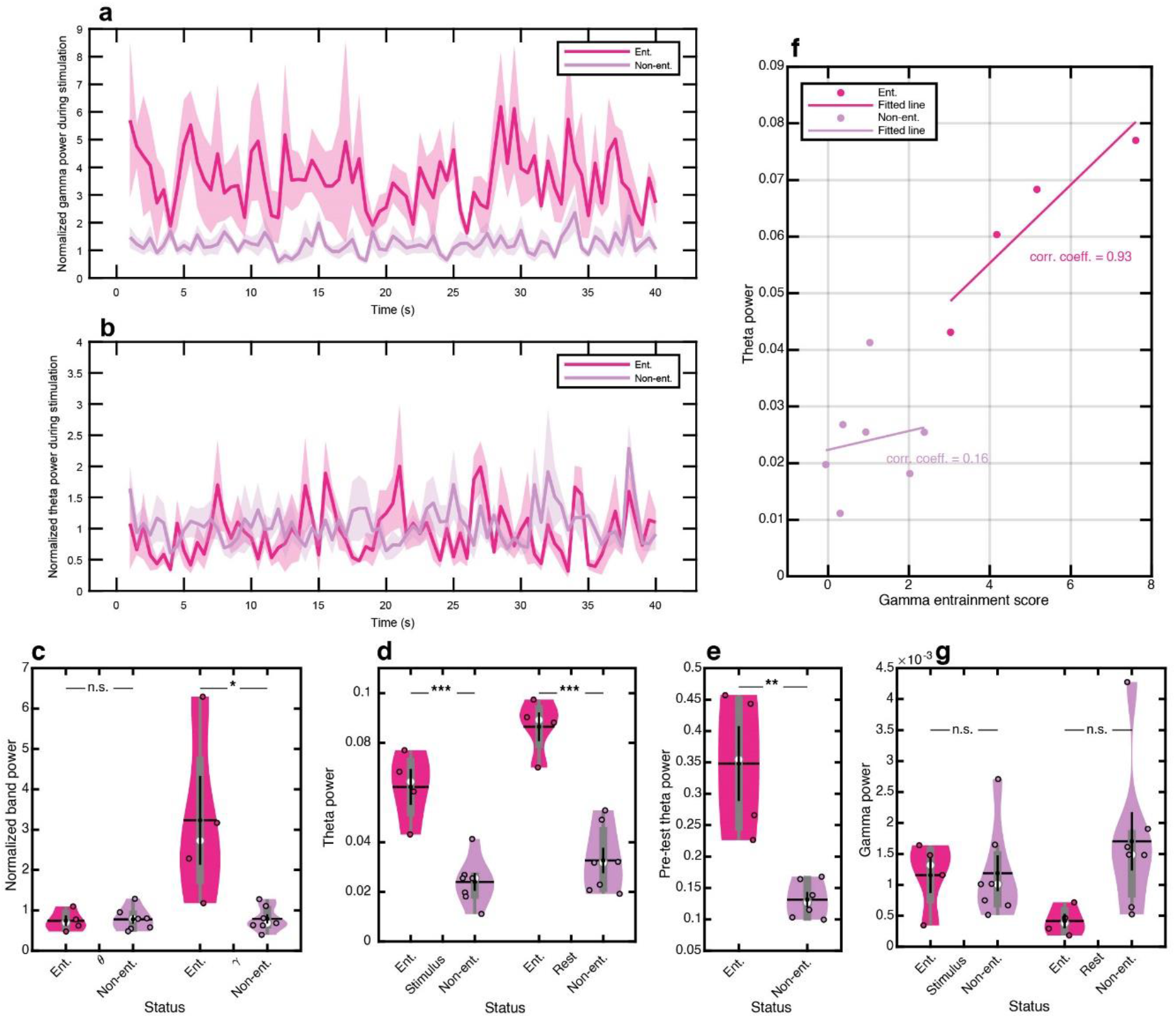
Theta and gamma band powers ensemble averaged in the stimulus and rest cycles for channel Fz. (**a**) Gamma power during stimulus cycles in 1sec sliding windows with 0.5sec overlap normalized to the total power of gamma in rest cycles for the entrained and non-entrained groups. (**b**) Same as **a** for theta power. (**c**) Total normalized (stimulus over rest) theta power (left) and gamma power (right) for both groups. (**d**) Theta power for stimulus (left) and rest (right) cycles for both groups. (**e**) Rest-state theta power for both groups calculated over 60sec before the start of the main task (rest-state data was not recorded from participant S9 in the non-entrained group). (**f**) Theta power versus gamma entrainment score. The entrained response is highly correlated with the theta power during stimulus cycles in the entrained group. This correlation is very low for the non-entrained group (Pearson correlation is used). (**g**) Same as **d** for the gamma power. Data expressed as mean ± SEM in shaded format in **a-b** and black lines in **c-e, g** as well as box-and-whisker plots in gray, violin plots representing the estimated normal distribution for each group, empty circles corresponding to each participant and white circles showing the median in the violin plots. *, p < 0.05; ***, p < 0.001. n.s.: not significant, Ent.: Entrained, Non-ent.: Non-entrained.

While normalized power can be used to determine the stimulation’s effect compared to rest, it cannot help assess the absolute power-differences between the two groups for the stimulation and rest cycles separately. Thus, we further analyze power spectra without normalization. Fig. 2d shows that the theta power is significantly higher for the entrained group in both stimulation (ttest2, t_9_ = 5.3826, p-value = 4.43e-04) and rest (ttest2, t_9_ = 6.6534, p-value = 9.34e-05) intervals. In other words, high theta power is present during the entire task in the entrained group. The concurrent presence of high theta power when gamma entrainment occurs can serve as a possible explanation for the superior performance of the entrained group in terms of the characteristics of the involved oscillatory bands. Remarkably, the presence of high theta power in the EEG signal recorded one minute before the start of the task can indeed serve as a predictor of the quality of the ensuing gamma entrainment (see Fig. 2e; ttest2, t_8_ = 4.3778, p-value = 0.0024). To further examine the role of theta power in facilitating gamma entrainment, we evaluate the correlation between them. As Fig. 2f indicates, not only higher theta power is present in the entrained group, it also registers higher values with higher entrainment scores. The relatively large positive correlation coefficient between the gamma entrainment score and the theta power (corr. coeff. = 0.93) suggests that the presence of more theta power may facilitate stronger gamma entrainment. Low values of the theta power and its rather flat relationship with the gamma entrainment score (corr. coeff. = 0.16) in the non-entrained group (Fig. 2f) provide further support for the effective role of theta oscillations in the quality of gamma entrainment.

According to Fig. 2g, the gamma power near the stimulation frequency is relatively lower for the entrained group during the rest cycles compared to the non-entrained group (ttest2, t_9_ = − 2.0061, p-value = 0.0758). It is noteworthy that there is an increase in the gamma power for the entrained group from the rest to stimulation cycles (ttest2, t_6_ = 2.3867, p-value = 0.0543), which reflects gamma entrainment in this group. On the other hand, a decrease in the gamma power is observed in the non-entrained group.

### Neural activity is highly synchronized during stimulation in the entrained group

The phase of the 40Hz frequency component provides additional information about the quality of gamma entrainment. By partitioning the recorded signals of channels Fz and Pz into one second windows, we extract the phase of the 40Hz component in each window and display these values as polar histograms for the stimulus and rest cycles. Figs. 3a-b show these histograms respectively for one entrained and one non-entrained participant as examples (see supplementary Figs. 2a-b for histograms of all participants). These plots suggest that for the entrained group, the distribution of the phases during the stimulation cycles is concentrated around a specific angle, whereas the responses of the non-entrained group show a large phase spread. The persistence observed for the phase of the 40Hz component during stimulation is a reflection of sustained synchronous activity of the entrained network. In other words, a constant phase across consecutive time windows means persistent delay, which is an indication of highly synchronized activity vis-à-vis the input driving signal. Additionally, the distributions of phase values in the stimulation cycles for the entrained group demonstrate about 180-degree differences between Fz and Pz (see supplementary Fig. 2a; rows 2 and 5). This phase difference offers further evidence for a persistent phase relationship in the neural activity across a large area of the brain. Moreover, it is noteworthy that activity with persistent phase occurs at all frontal, parietal, and occipital channels and notably at F3, Fz, F4, P3, Pz, P4, O1, and O2, where high entrainment score also exists (see supplementary Fig. 22). Remarkably, the central phase angle of the frontal channels (F3 and F4) follows the Fz phase whereas the parietal (P3 and P4) and occipital (O1 and O2) channels follow Pz with an around 180-degree disparity to the phase of the frontal nodes.

**Figure 3.**
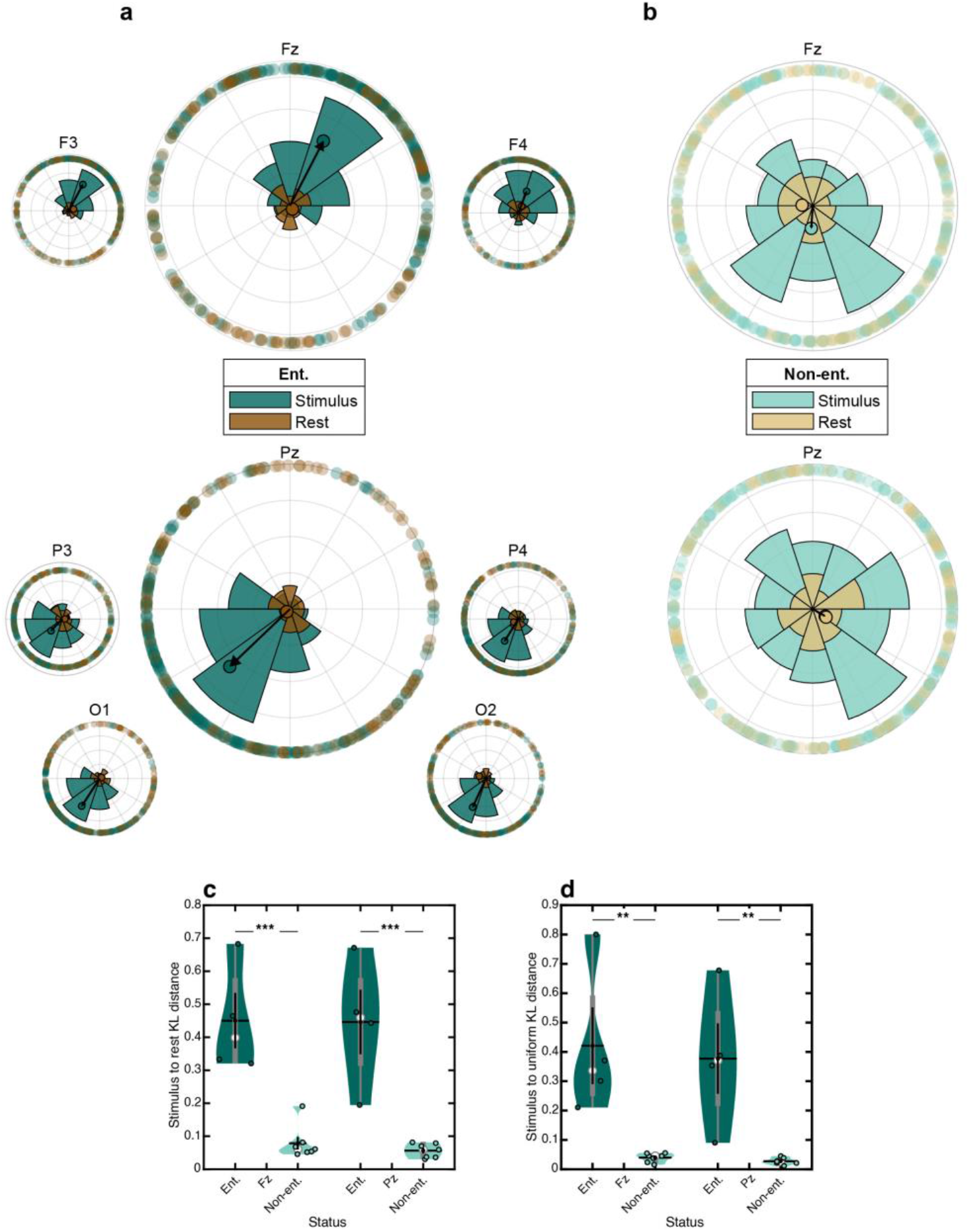
Neural activity is highly synchronized during stimulation in the entrained group. (**a**) Polar histograms of unit-amplitude phasors of the 40Hz component in 1sec sliding windows for the stimulus and rest cycles for a representative participant (S1) in the entrained group, recorded from the frontal (top) and parietal/occipital (bottom) channels. The colored circles on the perimeter show the phases of each 1sec sample. The empty circles represent average phasors of each 1sec record (for stimulus and rest sets). The black arrows correspond to the mean vector for the stimulus cycle set, with large values indicating highly concentrated response phases. Similar mean angles are observed for the frontal (F3, Fz, F4) channels. Also similar mean angles are observed for the parietal/occipital (P3, Pz, P4, O1, O2) channels with an approximate 180-degree disparity to the frontal mean angles. All these phenomena are observed in all participants in the entrained group (see supplementary Fig. 2a). (**b**) Same as **a** for a participant (S2) in the non-entrained group, showing results for Fz and Pz as representatives for the frontal and parietal/occipital regions, respectively. The phases of the response set during the stimulus cycles are spread out and hence the mean vectors have small amplitudes. (**c**) The KL distance between the stimulus and rest phase distributions shown in **a** and **b** averaged over all participants in each group. (**d**) Same as **c** but the KL distance is calculated between the stimulus phase distribution and a uniform distribution. Data expressed in **c-d** as mean ± SEM in black lines, box-and-whisker plots in gray, violin plots representing the estimated normal distribution for each group, empty circles corresponding to each participant and white circles showing the median. **, p < 0.01; ***, p < 0.001. KL distance: Kullback–Leibler distance, Ent.: Entrained, Non-ent.: Non-entrained.

On the contrary, in the non-entrained group both the stimulus and rest cycles produce phase values that are widely distributed across the entire range. This phase spread indicates that the oscillatory neural activity does not maintain a persistent phase across time, and that no synchronized activity is instilled in the network by the external stimulant.

To validate these deductions, we use the Kullback–Leibler (KL) divergence which measures the distance between two distributions. The KL distance of the phase distributions in the stimulus and rest cycles is significantly higher in the entrained group compared to the non-entrained group (ttest2, t_9_ = 5.6124, p-value = 3.2902e-04 for Fz, and t_9_ = 5.4459, p-value = 4.0784e-04 for Pz; Fig. 3c). This difference is an indication of a large disparity between the two groups in the brain’s synchronous activity caused by entrainment. We also measure the KL distance of the phase responses in the stimulus cycle to a uniformly distributed random set for the two groups, and the distance is again significantly greater for the entrained group compared to the non-entrained group (ttest2, t_9_ = 4.0147, p-value = 0.0030 for Fz, and t_9_ = 4.0135, p-value = 0.003 for Pz; Fig. 3d). These two observations together suggest that the entrained 40Hz neural activity due to the external auditory stimulant follows a persistent phase course in the entrained group whereas such temporal synchronization is not present in the non-entrained group (or during the rest cycle in the entrained group, further indicating the effective entrainability of this group).

### The entrained brain oscillations are spatially coupled

As discussed above and illustrated in Fig. 3a and supplementary Fig. 2a, the distributions of the response phases for Fz and Pz are persistently related to each other with a 180 degree shift during stimulation cycles for the entrained group. To assess whether the entrained gamma activity recorded from these two channels as well as other pairs of recorded channels maintain persistent synchronization, we calculate the phase locking value (PLV) for all channel pairs during the stimulation and rest cycles. PLV calculates how two oscillatory signals fluctuate congruently by temporal averaging of their phase-locking vectors. In our study, we use PLV as a measure to evaluate how well the gamma band activity due to entrainment is spatially coupled. We compute the PLV difference of the stimulus and rest responses in the gamma band for each participant. The results indicate a significant distinction between the entrained and non-entrained groups when PLV is measured between Fz and Pz (ttest2, t_9_ = 3.0303, p-value = 0.0142; Fig. 4a). Interestingly, most other parietal and occipital channels also show strong phase-locking values to frontal channels in the entrained group (Fig. 4b). High values of PLV for distant channels provide further evidence that the entrained 40Hz oscillations are spatially coupled in the entrained group.

**Figure 4.**
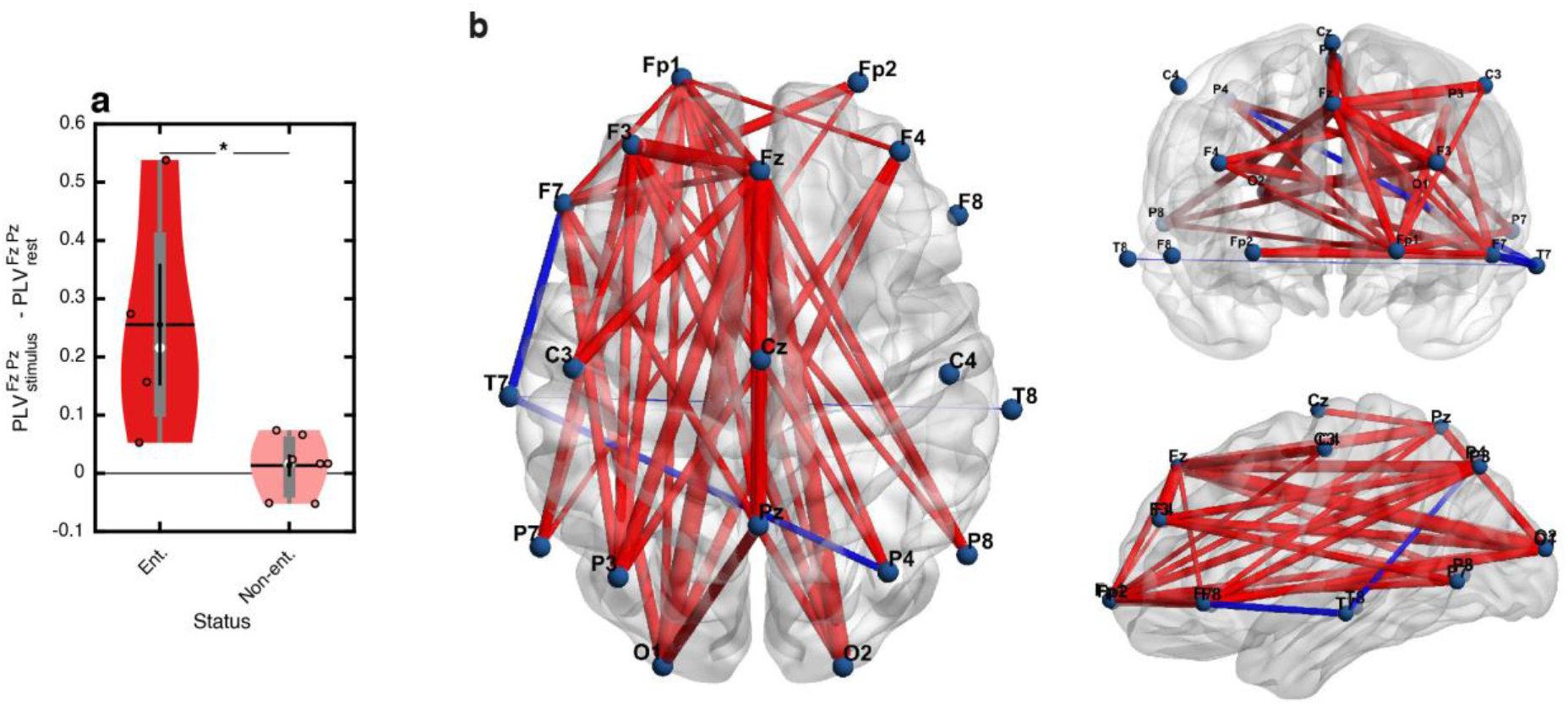
The entrained brain oscillations are spatially coupled. (**a**) The difference between the stimulus and rest responses in the phase locking value (PLV) measured between the Fz and Pz channels in the target gamma-band frequency is significantly larger for the entrained group compared to the non-entrained group. (**b**) Brain connectivity graph with edges between each pair of channels drawn based on the PLV difference of the stimulus and rest responses. Only the edge values that are significantly different between the two groups are shown. Edges are color coded with red indicating positive values (the PLV difference for the entrained group is larger than that for the non-entrained group) and blue indicating negative values, and thickness coded with the absolute value of the difference. Note the long-range connections from the frontal areas to the parietal/occipital areas depicted in axial (left), coronal (top right), and sagittal (bottom right) views. Data is expressed in **a** as mean ± SEM in black lines, box-and-whisker plots in gray, violin plots representing the estimated normal distribution for each group, empty circles corresponding to each participant and white circles showing the median. *, p < 0.05. Ent.: Entrained, Non-ent.: Non-entrained.

### The entrained gamma oscillations are highly theta coupled

Cross frequency coupling (CFC) exists in a healthy brain network^41,42^. Theta-gamma coupling is a phase-amplitude coupling (PAC) representing the modulation of the amplitude of gamma band activity by the phase of the theta band oscillations^42^. This modulation is reported to be involved in tasks related to working memory, spatial memory, and binding sensory data which all involve the hippocampus^13,43–45^. The quality of theta-gamma PAC is affected by dementia due to AD^14,15^. It is hence worth considering the effect of entrainment on the coupling. To compare how the coupling effect differs in stimulation and rest cycles for the two groups, we measure the mean vector length (MVL) value of the theta-gamma coupled vectors in the recorded signal. Comodulograms plotted based on this measure provide insight into the interaction of the theta and gamma oscillations. Fig. 5a depicts MVL comodulograms for one entrained and one non-entrained participants for channel Fz during stimulus and rest cycles (supplementary Figs. 3a-b illustrate MVL comodulograms for Fz for all participants). High MVL values at 40Hz during stimulation in the entrained participant in Fig. 5a (left) indicate that the amplitude of the entrained 40Hz oscillations is well modulated by the phase of theta oscillations. For the same participant, no noticeable coupling occurs during the rest cycles. No noticeable coupling is observed for the non-entrained participant during the stimulation or rest cycles. Across the entrained group, the difference of MVL values in the stimulus and rest cycles is significant (ttest2, t_9_ = 3.3359, p-value = 0.0087 for Fz, and t_9_ = 3.9255, p-value = 0.0035 for Pz; Fig. 5b. See supplementary Fig. 3c in which all significant channels are shown). An interesting point in the results of the non-entrained group evident from Fig. 5b is the lower mean value of the stimulus MVL compared to rest. In addition, the topographic distribution of MVL differences between the stimulus and rest cycles for the two groups is shown in Fig. 5c. Similar to the plots that show the entrainment strength (Fig. 1c) and the PLV measure (Fig. 4b), the strength of the theta-gamma PAC measured as MVL is also considerably higher on the frontal, parietal, and occipital channels. Moreover, the MVL value during stimulus cycles is significantly higher in the entrained group compared to the non-entrained group (ttest2, t_9_ = 3.5898, p-value = 0.0058 for Fz, and t_9_ = 2.4018, p-value = 0.0398 for Pz; Fig. 5d). These observations establish that effective coupling of theta-gamma activity occurs in the entrained group as a result of the 40Hz auditory stimulation.

**Figure 5.**
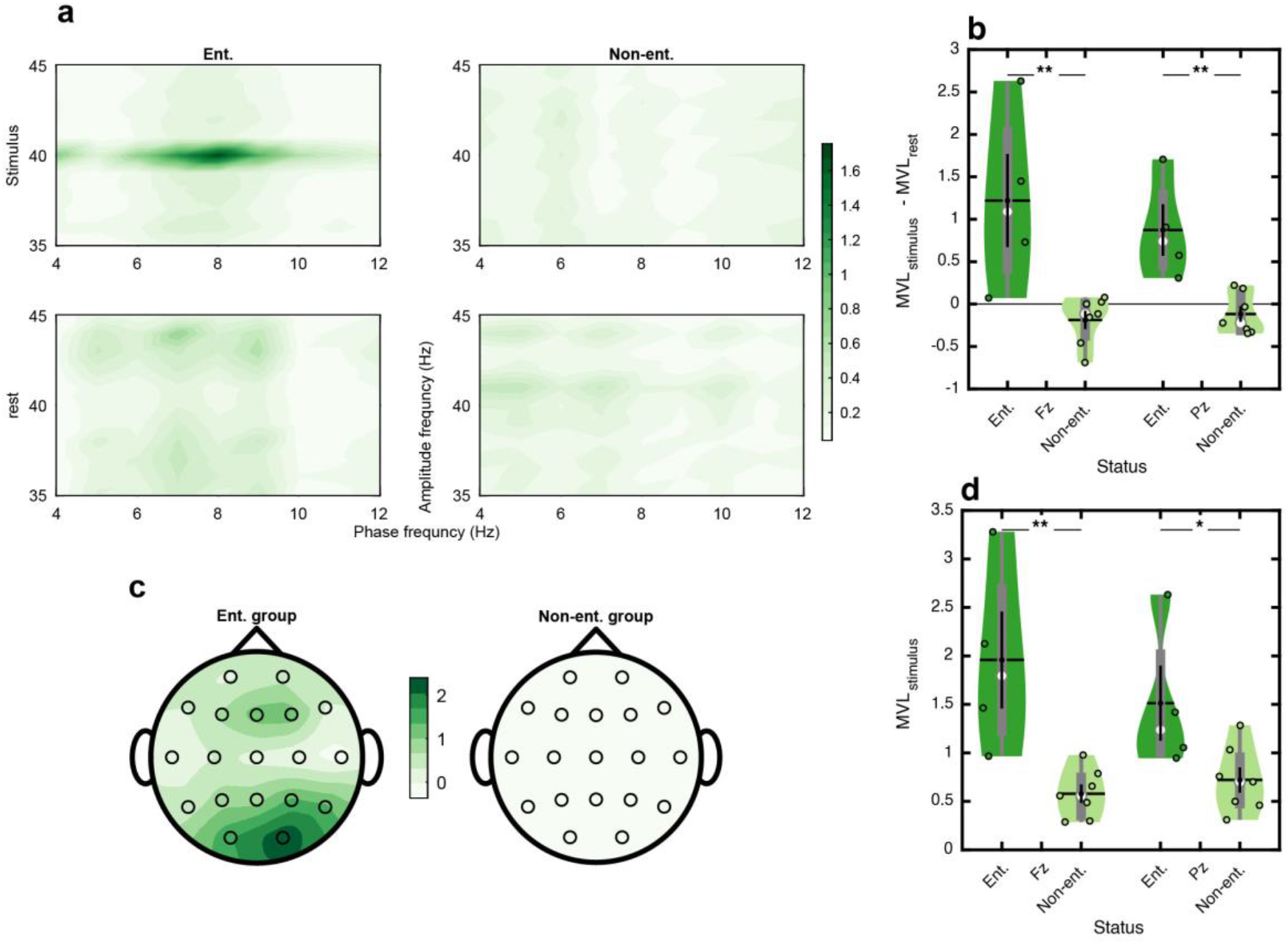
The entrained brain oscillations are highly theta-gamma coupled. (**a**) Comodulograms showing the strength of phase amplitude coupling (PAC) between theta band as the phase frequency and gamma band as the amplitude frequency oscillations based on the mean vector length (MVL) measurement during the stimulus (up) and rest (bottom) cycles for one representative participant (S1) in the entrained group (left) and one (S2) in the non-entrained group (right) (see supplementary Figs. 3a-b for similar plots for all participants). Note the high coupling value around the 40 Hz frequency component for the entrained participant due to the external stimulus. (**b**) Difference of MVL values in the stimulus and rest cycles for each group on the Fz and Pz channels. It is noteworthy that the MVL value is higher during rest than in the stimulus cycles for the non-entrained group (negative values for the mean MVL difference). (**c**) The topographic distribution of the MVL difference between the stimulus and rest cycles averaged for each group. In the entrained group, the frontal, parietal and occipital channels are highly theta-gamma coupled, which are also highly entrained (see the similarity of the patterns to the plots of Fig. 1c). (**d**) MVL values in the stimulus cycles for both groups measured for the Fz and Pz channels. Data expressed in **b**, **d** as mean ± SEM in black lines, box-and-whisker plots in gray, violin plots representing the estimated normal distribution for each group, empty circles corresponding to each participant and white circles showing the median. *, p < 0.05; **, p < 0.01. Ent.: Entrained, Non-ent.: Non-entrained.

## Discussion

Synchronized oscillations matter in cognitive functions and are manifested as temporal and spatial synchrony of oscillatory bands in different brain regions and intraregional phase-amplitude coupled activity^39,40^. The importance of synchronized activity increases considering the reported departure of its characteristics from normal levels due to dementia and AD^39,40^. Functions of the hippocampus in generating and coupling brain oscillations which even affect long-range communications in the brain have been under study^13,44^. While the generation of theta and gamma oscillations as the key rhythms in sensory data binding and memory processes has been an established function of the hippocamps^46^, the mechanisms involved in generating these oscillations and their interactions are still under examination. New studies have shown that reciprocal interactions of excitatory pyramidal cells (PCs) with inhibitory interneurons such as the PV cells and somatostatin (SST) contribute to the gamma activity of GABAergic cells^13,47^, the inhibitory tone of which plays a primary role in regulating network functions^4,5,47^. Moreover, malfunctions in inhibitory signaling or loss of inhibitory synapses can cause arrhythmic brain oscillations^1^. Theta oscillations also originate by the intrinsic activity of the hippocampus or due to the inhibition of PCs by the upstream area (medial septum) and play a role in modulating the gamma band amplitude^13,47^. A noteworthy observation in our study is that the entrained group shows high theta power during rest which accompanies low gamma activity (see Figs. 2d, g). In addition, an increase in the gamma power from the rest to stimulus cycles due to entrainment in this group is accompanied by a lowered theta power (see Figs. 2d, g). This counter-balancing effect may reflect the inhibition of the PC firing and hence reduced gamma power due to theta activity^13^.

### Entrainment and plasticity

New treatment approaches for AD can aim to rehash the affected neuronal communication pathways by resuscitating the weakened synapses through entraining the preexisting synaptic activity. Entraining the brain in repetitive sessions has been reported to hinder plaque formation^29^. A relevant question is how entrainment is able to enliven the characteristics reflecting the quality of synchronized oscillatory activity.

Neuronal populations engaged in synchronized activity resemble a group of musicians playing in an orchestra producing distinct rhythms, which they tune temporally and collectively by following a reference beat. As synaptic loss extends due to the progress of AD, the production of rhythmic oscillations loses synchronicity across the network as large neuronal populations turn into rather isolated smaller groups which cannot keep track of each other’s rhythms as they did before, leading to arrhythmic brain activity^4^. The entraining stimulant enters the scene as a new conductor (more precisely a metronome), pumping rhythmic beats into the neuronal populations and forcing oscillatory activity at the entrained frequency into synchronicity. Indicating factors for the efficacy of entrainment can hence be the temporal span and the spatial spread across which synchronized gamma activity is induced by the stimulant. Fig. 1c shows the rather large area in which high power activity is observed around the stimulant frequency. Furthermore, the entrained activity maintains a persistent phase across successive stimulation cycles as evidenced in Fig. 3a, indicating that the induced activity is temporally synchronized. Moreover, the spatial coupling of the induced gamma oscillations in the frontal and parietal/occipital areas points to the ability of the entrainment pump to bring different neural circuitry across the brain into synchrony at the target frequency. However, significantly lower levels of synchronization between these regions presents among the non-entrained group. Fig. 4b illustrates the distinction in spatial synchrony between the two groups in the form of a connectivity map, in which the strongest differentiating edges are predominantly between the mentioned areas. This spatial synchrony in the entrained brains mimics the characteristics of long-range communication activities in the healthy brain which are involved in sensory processing, memory and cognition^13,44,48^, while synchronous neural activity only exists locally in the non-entrained brains.

Finally, the induced gamma oscillations are coupled with the phase of theta activity as evidenced by high PAC values in comodulograms like Fig. 5a. This coupling follows a form of inter-band synchronization reported to occur in binding activities that involve the hippocampus^13,44,48^. The similarity of the spatial topographies of high gamma power (Fig. 1c) and high theta-gamma PAC (Fig. 5c) indicates that the entrained gamma oscillations tend to couple their amplitudes with the phase of the underlying theta activity present in the network.

How can the entrainment of gamma band oscillatory activity lead to the therapeutic effects reported for AD ^27–30,33,34^? One mechanism which may help explain these effects is the principle of neuroplasticity, which states that the synaptic links between neurons experiencing synchronized activity strengthen over time. Through enticing simultaneous neural activity, entrainment contributes to binding neural pathways and boosting of synaptic weights in neuronal populations that are forced to undergo synchronized activity.

### The role of theta oscillations in entrainment

There still remain other important questions. What is the role of theta oscillations in the process of entrainment when the stimulant frequency lies in the gamma band? Theta power plays a key role in human memory^48^. One observation in our study is that theta oscillations matter in entrainment as high theta power leads to more intensive entrainment (Fig. 2f). To justify this effect, we can consider the process of theta-nested gamma oscillations which reflects the interaction of PCs and interneurons. This interaction involves feedback loops between excitatory and inhibitory populations in the hippocampus, through which PCs induce theta cycles in the interneurons, which in turn send inhibitory signals to the PCs as gamma cycles. Theta-gamma coupled oscillations can thus be described by a pyramidal interneuronal network gamma (PING) model^49^, which describes the interaction of the theta and gamma oscillatory activities. One arresting idea triggered by the cyclic nature of theta-gamma interactions suggested by the PING model is that both gamma stimulations^23^, as in our study, and theta stimulations^47,49–52^ be employed to increase the coupling strength and even the gamma power. More generally, the premise of the PING model motivates that any stimulating intervention in the excitatory-inhibitory feedback loop can be used to trigger the cyclic circuit no matter if the stimulant is in the theta or gamma bands. The loop’s circular property ensures that starting from one point in the loop leads to traversing the cycle and triggering the excitatory and inhibitory activities along the way. This line of reasoning suggests that theta entrainment may lead to similar therapeutic effects reported for gamma entrainment. While such variations of the entrainment process deserve further examination, a few initial observations can be derived from the current study which point to using the gamma band as the preferred stimulus. First, gamma entrainment appears to be a more effective trigger since the high theta power present in the entrained group during the rest cycles did not autonomously lead to gamma generation and coupling. Second, the hippocampus acts as a gamma generator^50^, and since the gamma quality is affected by AD^4^, direct gamma entrainment may prove more effective in repairing the cycle. Recent studies have reported improvements on patients using gamma entrainment^30,34^. In a study based on stimulating PV cells of the hippocampus invasively via optogenetics in a mouse model of AD, rehabilitated theta-gamma coupling and gamma power were reported along with better performance in an object and place recognition task^23^. Our results provide evidence for significantly improved coupling performance due to entrainment in humans, and demonstrates that this effect can be measured based on EEG data.

These points aside, the role of the theta-band activity should not be overlooked in entrainment studies as there are evidences on improving performance in memory tasks due to theta entrainment^53^. Our study led to remarkable observations on the role of theta power in gamma entrainment, namely its very high correlation with the gamma entrainment score (Fig. 2f), and the efficacy of the rest-state theta power in predicting successful entrainment (Fig. 2e). These observations elude to the idea that the presence of a theta oscillatory component in the stimulant might improve the quality of gamma entrainment in non-entrained participants through increasing theta activity in the brain. Synthetic auditory stimulants can be created in different ways to contain both gamma and theta oscillations. In a recent report^54^, we used chirp segments of natural canary song, which contains some level of intrinsic theta-gamma coupling, and observed high-quality brain entrainment for a group of participants.

### Multimodal entrainment

Another area for further examination is the use of different sensory modalities in entraining the brain. Auditory and visual modalities alone have been shown to offer beneficial effects. Visual stimulation reduced the plaque level and tau phosphorylation in the visual cortex, whereas auditory stimulant did the same in CA1 as well as in the auditory cortex^29^. Interestingly, simultaneous audio-visual stimulation cleared even more regions including prefrontal cortices when employed in a similar stimulation time course^29^. As a result, the therapeutic effect of these modalities can be ordered from least to most as visual, auditory, and audio-visual. To assess whether this order of efficacy can be also revealed by the theta-gamma coupling strength, we separately applied visual, auditory, and audio-visual stimulation to a healthy individual and produced MVL comodulograms for all three cases (see Fig. 6). The audio-visual entrainment shows the most intensive coupling at the 40Hz frequency. Similar observations on enhanced theta-gamma coupling in presence of multimodal stimulation were reported in a recent study by our lab on a group of participants^55^. An interesting issue to further examine is whether entraining wider areas of the brain using multiple modalities can yield better therapeutic results through restoring long-range communication which is progressively impaired as AD affects larger areas of the brain^56^. Hence, employing other sensory modalities such as somatosensory or olfactory in addition to audio-visual entrainment may offer additional therapeutic value and can be the subject of further investigation.

**Figure 6.**
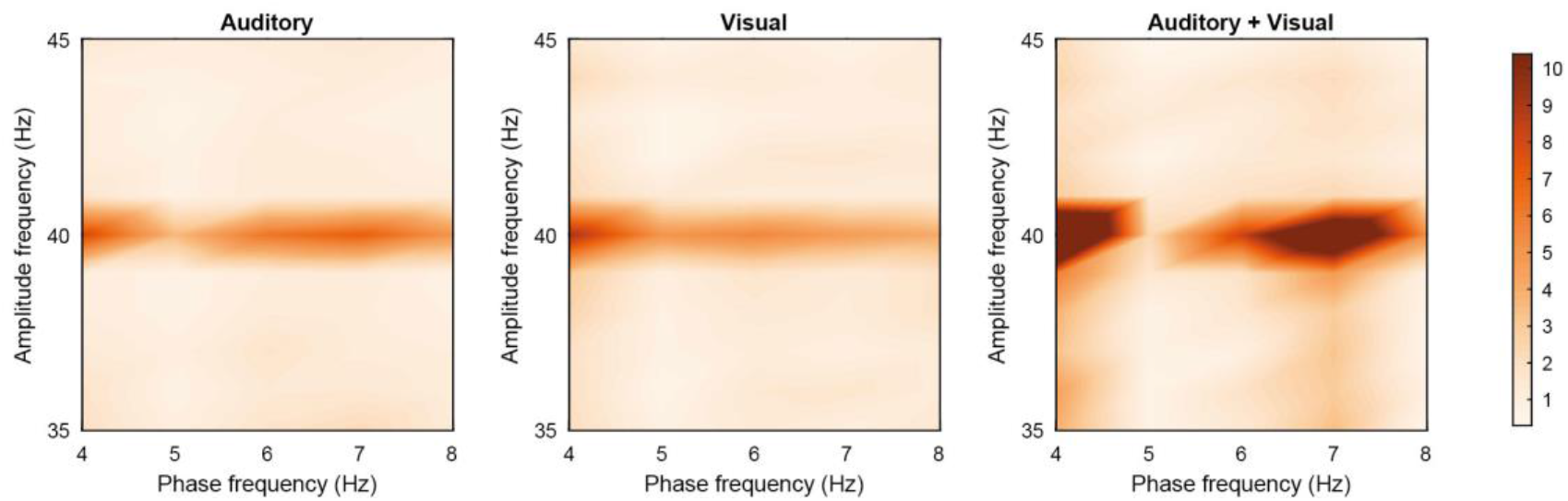
Multi-modal entrainment improves the phase amplitude coupling (PAC) of the theta and gamma oscillations. MVL comodulograms of the theta-gamma PAC for channel Fz for a healthy individual measured in three distinct tasks: auditory (left), visual (middle), and simultaneous auditory-visual (right), all with 40Hz entrainment.

### Limitations of our study

Our study had limitations in its scope and implementation in several ways. First, since our main objective has been to provide interpretive evidence for the therapeutic effects of gamma entrainment, we limited the scope of our data collection to only one session with multiple trials. This choice was to alleviate the burden of multiple visits to the clinic for our elderly participants and to minimize their risk of exposure to COVID-19 infection. While processing data acquired from one session already provided a multi-faceted interpretation for the mechanisms by which gamma entrainment improves network performance, obviously a longitudinal study is required to draw conclusions on how these network characteristics evolve over time.

Second, the duration of the stimulation and rest cycles within one session may affect the entrainment results. Furthermore, the effective cycle duration may be different for each participant and need customized tuning. Our study did not include such tuning to avoid elongated sessions for our participants, which might have been exhausting to them as well as causing distraction and diminished attention to the stimulant. Instead, we separately applied different stimulus and rest cycle durations on a group of young and healthy volunteers and based on the results of that study selected the intervals used in the current study. Cycle durations and session length do play a role in the entrainment quality and deserve further examination. It is likely that some of the participants whose response did not surpass our definition of entrainment could produce better responses with longer stimulation cycles. When feasible, personalized tuning of these attributes of the entrainment session can be the subject of extended work.

Third, and as initial results of other studies based on longitudinal assessment of the entrainment-based therapy on humans are being reported^35^, important practical questions such as at what stage of dementia or AD would gamma entrainment still be able to help reverse the symptoms of the disease, or how long will the improvements linger after the end of the therapy still remain outstanding. These questions also pertain to our approach of providing a network-based interpretation of the effects of entrainment, and deserve attention in further related studies.

## Methods

### Participants

Thirteen volunteers (five females, 57-89 years of age) were recruited from referrals to the memory clinic of Ziaeian Hospital in Tehran with memory performance complaints. A neurophysiologist from the Department of Geriatric Medicine of Ziaeian Hospital conducted all clinical procedures for this study. The participants’ age, level of education, preferred hand, and smoking history were recorded. Cognitive status was quantified using the mini-mental state examination (MMSE). A neurologist examined the probable AD state in each participant according to the latest guideline of the NIA-AA^57^ and performed the functional assessment scales test (FAST) on the participants.

This study was approved by the Review Board of Tehran University of Medical Science (Approval ID: IR.TUMS.MEDICINE.REC.1398.524) and all participants provided informed consent before participating and were free to withdraw at any time. Participant demographics, including age and neuropsychological scores, are outlined in Table 1.

**Table 1.** General information of all participants

| Index | Participant code | MMSE score | Gender | Age | Dementia state | Inclusion | Entrainment status |
| --- | --- | --- | --- | --- | --- | --- | --- |
| 1 | S1 | 28 | Male | 65 | Non-AD | Yes | Entrained |
| 2 | S2 | 23 | Female | 70 | Mild AD | Yes | Non- Entrained |
| 3 | S3 | 27 | Male | 75 | Non-AD | Yes | Entrained |
| 4 | S4 | 21 | Male | 82 | Mild AD | Yes | Non- Entrained |
| 5 | S5 | 21 | Male | 75 | Mild AD | Yes | Entrained |
| 6 | S6 | - | Female | 69 | - | No | - |
| 7 | S7 | 19 | Male | 89 | Mild AD | Yes | Non- Entrained |
| 8 | S8 | 24 | Male | 70 | Non-AD | Yes | Entrained |
| 9 | S9 | 26 | Female | 57 | Non-AD | Yes | Non- Entrained |
| 10 | S10 | 28 | Male | 81 | Non-AD | Yes | Non- Entrained |
| 11 | S11 | 20 | Female | 88 | Mild AD | Yes | Non- Entrained |
| 12 | S12 | 30 | Female | 63 | Non-AD | Yes | Non- Entrained |
| 13 | S13 | - | Male | 60 | - | No | - |

General exclusion criteria were: a history of stroke, traumatic brain injury, schizophrenia, major depressive disorders and electroconvulsive therapy (ECT) over the prior six months, or other neurodegenerative diseases (Parkinson’s disease, multi-system atrophy, cortico-basal degeneration, progressive supranuclear palsy). Two participants (S6 and S13) were excluded from the study as the diagnosis of their status required further examination not scoped in this study.

### EEG recording and preprocessing

All EEG data were recorded using 19 monopolar channels in the standard 10/20 system referenced to the earlobes, sampled at 250Hz, and the impedance of the electrodes was kept under 20kΩ. During the experiment, participants were seated comfortably with open eyes in a quiet room, and they were instructed to relax their body to avoid muscle artifacts and to move their head as little as possible. Before the main task, a one-minute data was recorded with open eyes for measuring raw resting-state potentials.

Data from all the participants were preprocessed identically following Makoto’s preprocessing pipeline^58^: Highpass filtering above 1Hz; removal of the line noise; rejecting potential bad channels; interpolating rejected channels; re-referencing data to the average; artifact subspace reconstruction (ASR); re-referencing data to the average again; estimating the brain source activity using independent component analysis (ICA); dipole fitting; rejecting bad dipoles (sources) for further cleaning the data. These preprocessing steps were performed using EEGLab^59^ toolbox in MATLAB. Finally, the data (each channel) were normalized to zero mean and unit variance for further analysis and making comparison among participants.

### Auditory stimulation

Two speakers were placed in front of the participant 50cm apart from each other and directly pointed at the participant’s ears at a distance of 50cm. The sound intensity was around −40dB within a fixed range for all participants. Before starting the task, the participant was asked if the volume was loud enough and the sound volume was set at a comfortable level for each participant. The auditory stimulus was a 5kHz carrier tone amplitude modulated with a 40Hz rectangular wave (40Hz On and Off cycles). Since a 40Hz tone cannot be easily heard, the 5KHz carrier frequency was used to render the 40Hz pulse train audible. In order to minimize the effect of the carrier sound, the duty cycle of the modulating 40Hz waveform was set to 4% (1ms of the 25ms cycle was On). The auditory stimulant was generated in MATLAB and played as a .wav file. This file consisted of 6 trials of 40sec stimulus interleaved by 20sec of rest (silence). Thus the data collected from each participant was a 340sec (6×40+5×20) EEG signal.

### Visual stimulation

To conduct an initial study on the effect of adding visual stimulation to the auditory stimulation, we conducted three sessions with 40sec of auditory, or visual, or audio-visual stimulation on a healthy volunteer (23 years old male) and recorded EEG data during all three sessions. The visual stimulant was a 20Hz flickering white light produced by an array of LEDs and reflected from a white wall at 50cm distance in front of the participant (open eyes) with 50% On cycles (duty cycle = 50%). The data of the first 20sec of the stimulation cycles in each session were used for producing the MVL comodulograms. Due to the presence of harmonic frequencies in its pulse train^37^, the 20Hz stimulant can drive 40Hz oscillations in the brain.

### Definition of entrainment

We first decide on the occurrence of entrainment in each trial of each participant if the peak frequency amplitude of the response at the 40Hz stimulant frequency is at least three times the standard deviation away from the mean of response amplitudes in a range of adjacent frequencies (38Hz to 42Hz). In other words, if the z-score of the amplitude at 40Hz in the response compared to the responses at the local frequency neighborhood exceeds 3 (z-score > 3), the trial is considered entrained (see supplementary Fig. 1). We did this for each channel, trial, and participant.

Judging entrainment by EEG signals can be tricky due to the effect of the background activity of the brain which may overshadow the response to the main task. So to determine whether a channel was entrained for a participant, if the majority of trials (4 trials out of 6) were entrained according to the defined criterion, the channel was marked to be entrained.

### Gamma entrainment score

For quantitative analysis we define a parameter to describe how well a channel is entrained for the entire task. The z-score values of all six stimulation trials are averaged for each channel and the result is defined as the gamma entrainment score. This parameter significantly differs between the two entrained and non-entrained groups across most channels in the frontal, parietal and occipital lobes. We used the gamma entrainment score of channel Fz to divide the participants into entrained (n=4) and non-entrained (n=7) groups (supplementary Fig. 1b, ttest2, t_9_ = 4.7298, p-value = 0.0011 for Fz).

### Theta/gamma power

For power analysis, we averaged all six stimulation trials (each had a duration of 40s) as well as five rest trials (each had a duration of 20s) for each participant. Theta and gamma oscillations were then extracted by bandpass filtering the signals in the theta bandwidth (4-8Hz) and the target gamma bandwidth (39-41Hz). For obtaining power time series, theta and gamma powers were calculated in 1sec sliding windows with 0.5sec overlap during stimulation and were normalized to the total rest power. For Fig. 2c, the total stimulus power in each band was normalized to the total rest power in that band. However, for Figs. 2d, g, the total power in each band and each state (rest and stimulus) was calculated separately without normalization. The theta power used in Fig. 2e is the total theta power calculated from the 60sec rest signal recorded before the main task. In Fig. 2f, we used linear curve-fitting on the average z-score values (gamma entrainment score) as the measure for entrainment strength and the total theta power during stimulus cycles. Correlation coefficients between these two variables were calculated following the Pearson definition.

### Kullback–Leibler (KL) divergence

We extracted the 40Hz Fourier phase of 1sec sliding windows (with no overlaps) through the entire 340sec task block for the Fz and Pz channels. The phase values of the time windows associated with the stimulus and rest cycles were separately used to create the polar histograms shown in Figs. 3a-b and supplementary Figs. 2a-b. Bin sizes of 36° were used to divide the entire phase range into 10 bins. By dividing each bin’s count by the total number of samples, we calculated the stimulus (P) and rest (Q) distributions of the 40Hz Fourier phase.

For measuring the difference between the two distributions *P*(*x*) and *Q*(*x*), we used KL divergence as follows:

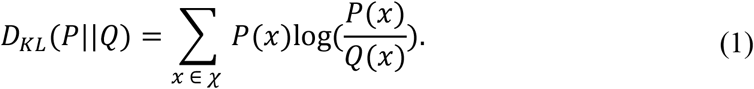

For the distance of a distribution *P*(*x*) from a uniform distribution *U*, this equation simplifies to:

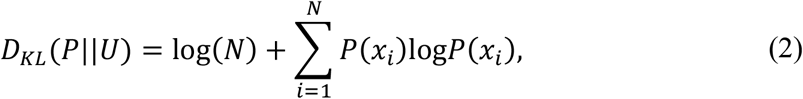

where N is the number of bins.

### Phase locking value (PLV)

After applying a narrow bandpass filter near 40Hz (39-41Hz) to the ensemble-averaged stimulus and rest signals, the phase of gamma oscillations was obtained by performing the Hilbert transform. To assess the phase synchronization between two distinct channels (Fz and Pz as an example), we averaged the Fz-Pz phase-locking vectors of all temporal samples. Defining *ϕ*_*Fz*_(*t*) and *ϕ*_*Pz*_(*t*) as the phases of gamma oscillations at the Fz and Pz channels, respectively, the PLV was computed as:

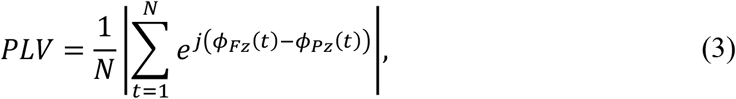

where *N* is the total number of temporal samples.

We calculated the PLV for both the stimulus and rest cycles and used the difference as a statistical variable. We only used the first 20sec of stimulus cycles in order to have the same number of temporal samples in both the stimulus and rest responses. The connectivity graphs showing the significant PLV differences between the stimulus and rest responses in the two groups were plotted using BrainNet Viewer^60^.

### Phase-amplitude coupling (PAC)

We calculated phase-amplitude coupling over windows of equal length in time to reach a fair comparison between the stimulus and rest trials. Thus, we chose the first 20sec of the stimulus cycles as well as full 20sec windows of the rest cycles after merging all trials to one stimulus and one rest by ensemble averaging each set of trials. Then, with the use of the mean vector length (MVL) method, we calculated the strength of theta-gamma coupling. The MVL factor is among the best measures for describing the phase-amplitude coupling effect, and can calculate the coupling strength between two frequency ranges or two distinct single frequencies^61,62^.

The envelope of the high-frequency amplitude component, 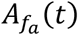, and the low-frequency phase component, 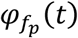, are extracted by the RID-Rihaczek distribution, which defines a signal’s complex energy distribution in the time-frequency domain (*C*_*t,f*_)^62^ leading to the time-frequency mean vector length (tf-MVL) measurement as follows:

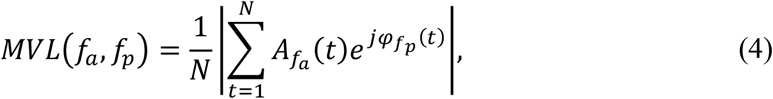

where *N* is the number of temporal samples. This equation represents the MVL for two distinct single frequencies (*f*_*a*_, *f*_*p*_) chosen within the gamma and theta bandwidths, respectively. To illustrate the phase-amplitude coupling of the two oscillatory bands, we plotted comodulograms, which are 2D plots that specify the MVL between different pairs of frequencies with the low frequency (*f*_*p*_) along the x-axis and the high frequency (*f*_*a*_) along the y-axis, and considering a frequency step size of 1Hz for each.

To indicate the coupling strength between the two oscillatory bands with a single number, first, we integrated the signal’s complex energy distribution in the time-frequency domain (*C*_*t,f*_) over the theta (*C*_*theta*(*t*)_) and gamma (*C*_*gamma*(*t*)_) bandwidths. Then, we used the Equation set (5) to extract the gamma amplitude time series, *A*_*gamma*_(*t*), in the range of 39-41Hz and the theta phase time series, *φ*_*theta*_ (*t*), in the range of 4-8Hz, and defined the coupling measurement for the two oscillatory ranges as in Equation (6).

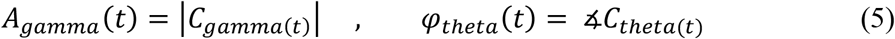

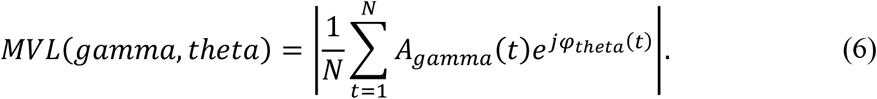

### Statistical analyses

All data are presented as mean ± standard error mean 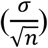 in black lines, and as box-and-whisker plots in gray, as well as violin plots representing the estimated normal distribution of samples in each group. Moreover, the samples are shown as empty circles and the median as white circle. All tests of significance are two-sided t-tests and are based on variance equality. Test details are described where applicable. p < 0.05 was considered statistically significant. *, p < 0.05; **, p < 0.01; ***, p < 0.001, ****, p < 0.0001.

## Acknowledgments

The authors wish to thank Ziaeian Hospital in Tehran for providing staff time and equipment for data collection in this study. We are grateful to the patients and their families who participated in this study. This work was partially funded by the Cognitive Sciences & Technologies Council of Iran and by the grant G970736 from Sharif University of Technology, which covered the cost of data collection. The funders had no role in the study conceptualization and design, data collection and analysis, decision to publish, or preparation of the manuscript.

## Author contributions

HA conceptualized the work. ML and HA designed the experiments. ML conducted the study. ML and ZV collected the data. ML and AA analyzed the data. ML, AA, and HA wrote the manuscript. HA and ZV supervised the work. All authors edited and approved the manuscript.

## Data availability

The datasets generated and analyzed in the current study are available at the OpenNeuro repository.

The data used for the main study is available at: 10.18112/openneuro.ds003800.v1.0.0

The data of the multi-sensory experiment is available at: 10.18112/openneuro.ds003805.v1.0.0

## Code availability

The codes for generating the results presented in the manuscript are available at: https://github.com/m-l-3/Gamma-Entrainment

