## Supplementary Materials for "Gamma Entrainment Improves Synchronization Deficits in Dementia Patients"

### Statistics

**Sample size**

The aim of this study has been to investigate the effect of gamma entrainment on demented patients. We expected to find statistical differences between the participants suffering from mild AD and those in the earlier stages of dementia due to normal ageing, and hence used the MMSE score as the differentiating parameter for sample size calculation.

Following equation (1) for calculating the sample size[1]:

$$n = \frac{2(Z_{\alpha} + Z_{1-\beta})^2 \sigma^2}{\Delta^2}, \quad (1)$$

we set $Z_{\alpha}$ to 1.96 for having a 5% significance (two-sided) and $Z_{1-\beta}$ to 0.8416 for a power of 80%. Reports published on associating the MMSE score with the status of dementia suggest a score of 20 to 24 for mild dementia[2]. So we expected a mean score of 22 for the mild AD patients and a mean score of 27 for normal ageing, yielding the mean difference value of $\Delta = 5$. In addition, we set the standard deviation ($\sigma$) of both scores to 3 according to previous studies conducted in our lab[3]. Using these values in equation (1) yields an estimated sample size of 6 per group.

As the current study was conducted, we further monitored the gamma entrainment score as a key parameter associated with the quality of entrainment, and employed it to divide the participants into two groups of entrained and non-entrained. This allowed us to assess the adequacy of the existing participant number in yielding significant results. Adhering to the principle of mitigating burden and risk (including the risk of exposure to coronavirus) for the target elderly population, we paused the data collection process once the parameters aimed to be evaluated in the study (see Methods) showed significant differences between the two groups of tested participants. Then, we re-evaluated the sample size using the statistics of the gamma entrainment score. According to Supplementary Fig. 2b, the estimated effect size ($\Delta$) in this case is approximately 4 and the standard deviation ($\sigma$) is around 2, yielding a sample size of 4 per group.

[1] Department of Electrical Engineering, Sharif University of Technology, Tehran, Iran
[2] Department of Geriatric Medicine, Ziaeian Hospital, Tehran University of Medical Sciences, Tehran, Iran

#### Data exclusions

Based on the exclusion criteria (see Methods), two participants whose conditions needed further examination were excluded from the study.

#### Replication

As our study aimed to assess the effect of a procedure reported in other studies from a new perspective, replication of the procedures was not relevant to our study.

#### Randomization

The aim of our study was to evaluate the network-based effects of a reported procedure involving the entrainment of brain oscillations. Therefore, the procedure was conducted on all participants and hence randomization of participants was not relevant to our study.

#### Blinding

The data analysts were completely blinded to the clinical state and demographic information of the participants, as well as their MMSE and other psychophysical scores and dementia state. In addition, EEG technicians were also blinded to the cognitive state of the participants and their MMSE scores as well as the definition and values of the parameters analyzed in the study.

### Supplementary figures

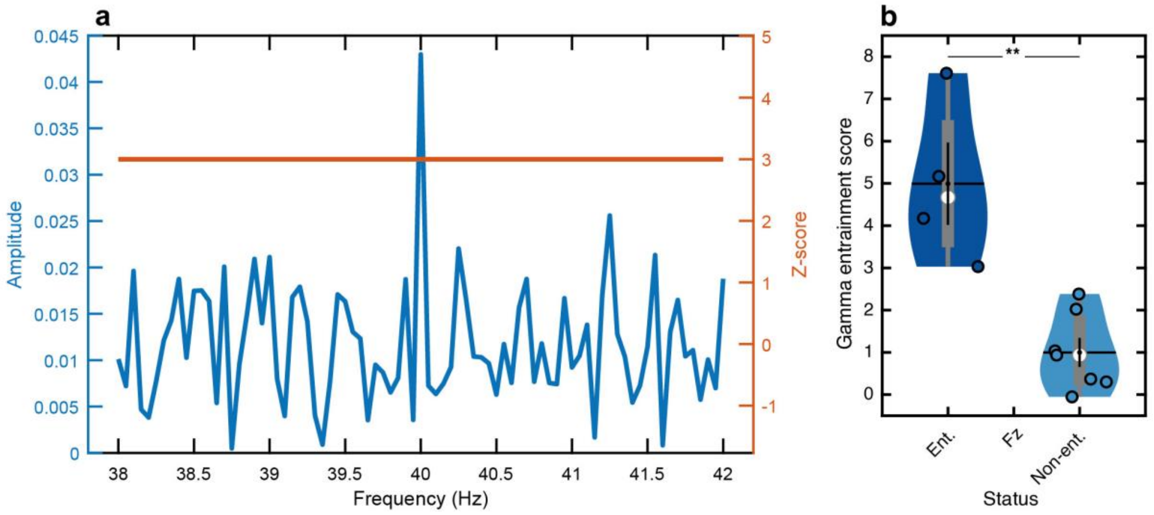


**Supplementary figure 1** Definition of entrainment in the recorded response and the method of grouping the participants. (**a**) The occurrence of entrainment in a trial is decided if a distinct peak is observed at the 40Hz component in the frequency response of the recorded signal with an amplitude at least three times the standard deviation away from the mean of response amplitudes in a range of adjacent frequencies (z-score value > 3). (**b**) The statistics of the gamma entrainment score on the Fz channel for the entrained and non-entrained groups (mean ± SEM in black lines, box-and-whisker plots in gray, violin plots representing the estimated normal distribution, empty circles corresponding to each participant and white circles showing the median). The trial-averaged z-score value of the 40Hz component is defined as the gamma entrainment score. This parameter shows how each channel is entrained in the entire task. Based on the gamma entrainment score of the Fz channel, the participants are grouped into two significantly distinct groups of entrained (n=4) and non-entrained (n=7). **, $p < 0.01$. Ent.: Entrained, Non-ent.: Non-entrained.

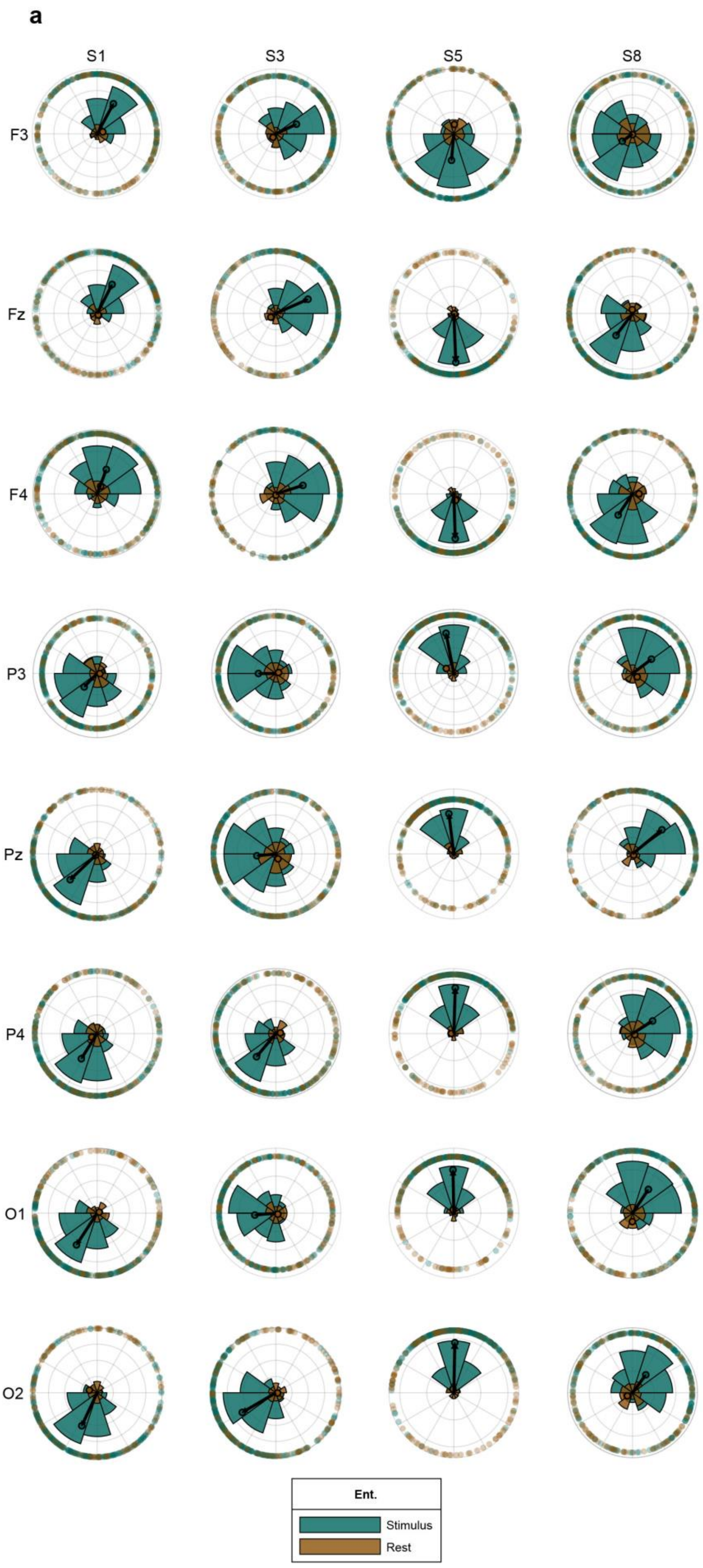
a
S1
S3
S5
S8
F3
Fz
F4
P3
Pz
P4
O1
O2
Ent.
Stimulus
Rest

**b**

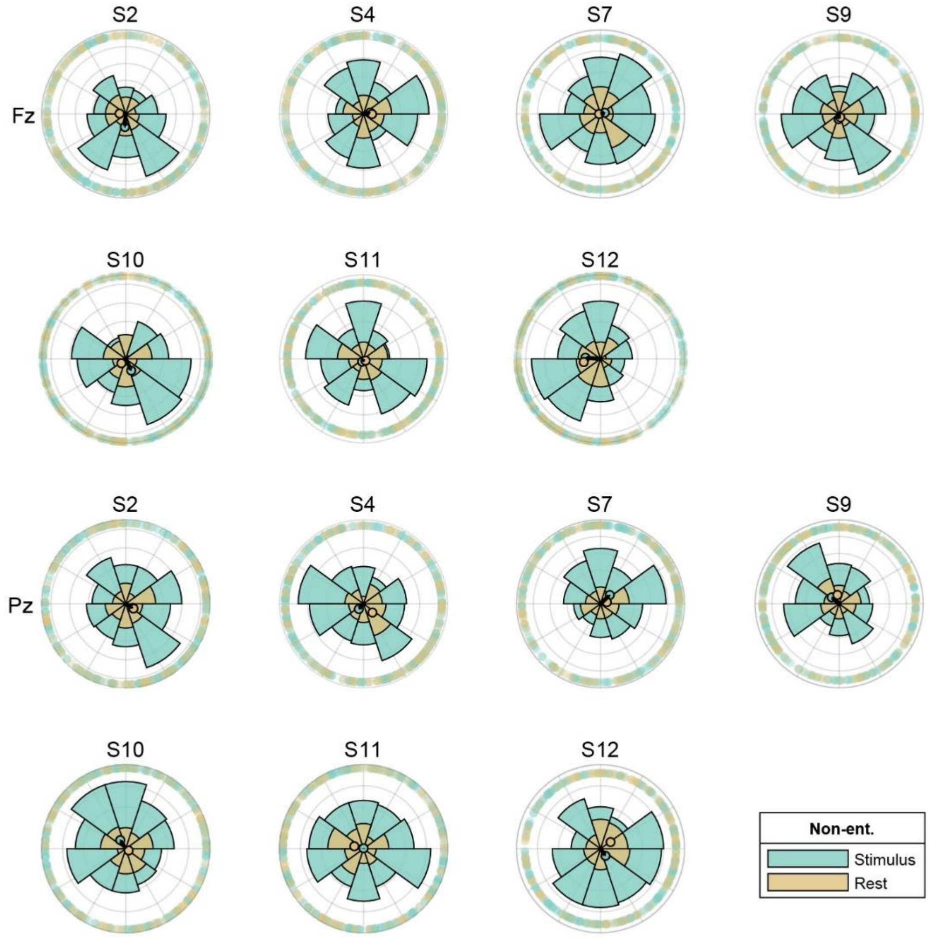
S2
S4
S7
S9
Fz
S10
S11
S12
S2
S4
S7
S9
Pz
S10
S11
S12
Non-ent.
Stimulus
Rest

**c**

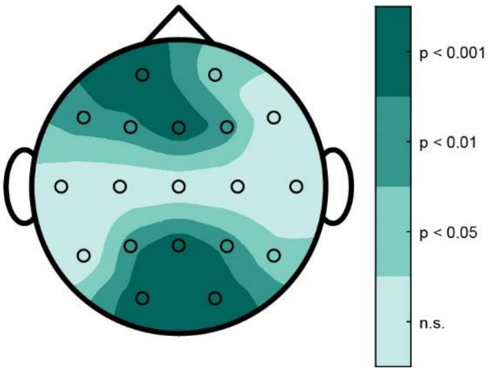
p < 0.001
p < 0.01
p < 0.05
n.s.

**Supplementary figure 2** Phase consistency of the 40Hz component during the entire task. (**a**) Polar histograms of unit-amplitude phasors of the 40Hz component in 1sec sliding windows for the stimulus and rest cycles for all participants in the entrained group (n = 4) for the frontal (F3, Fz, F4), parietal (P3, Pz, P4), and occipital (O1, O2) channels. The colored circles on the perimeter show the phases of each 1sec sample. The empty circles represent average phasors of each 1sec record. The black arrows correspond to the mean vector for the stimulus cycle set, with large values indicating that the response phases are highly concentrated. Note the similar mean angles for the frontal (F3, Fz, F4) channels and similar mean angles for the parietal/occipital (P3, Pz, P4, O1, O2) channels with an around 180-degree disparity to the phase of the frontal channels. This property also exists for more channels but for simplicity only the channels with highest entrainment scores (according to Fig. 1c) are shown. (**b**) Same as **a** for all participants in the non-entrained group (n = 7), depicting results for Fz (top) and Pz (bottom) as representatives for the frontal and parietal/occipital regions, respectively. Unlike the histograms in **a**, the phases of the response sets during the stimulus cycles are spread out and hence the mean vectors have small amplitudes. (**c**) Channels with significant stimulus to rest KL distance between the two groups. The p-value of the test of significance is color coded. Note the similarity with Fig. 1c, which is an indication of the co-occurrence of high entrainment performance and phase consistency. n.s.: not significant, Ent.: Entrained, Non-ent.: Non-entrained.

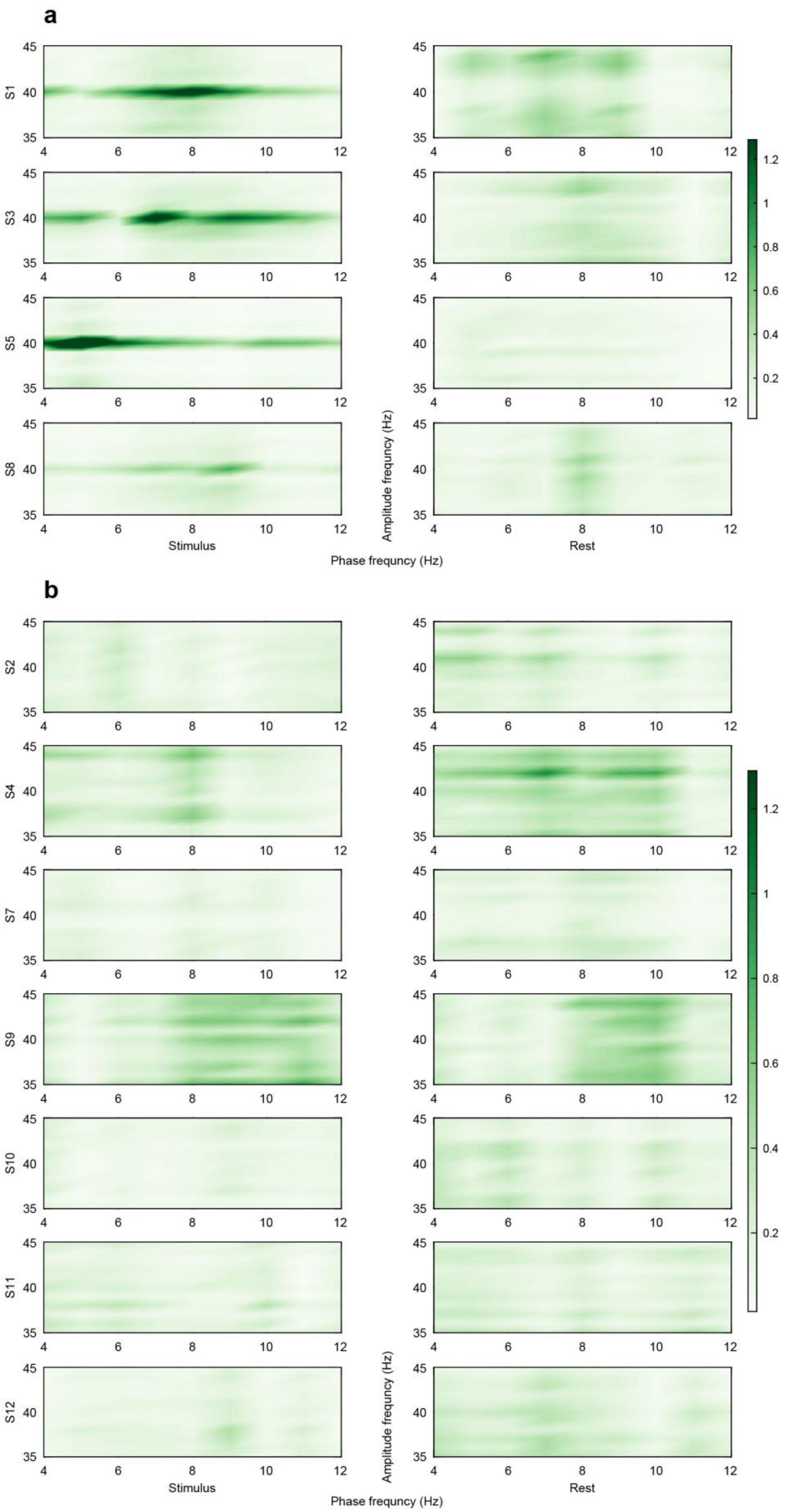
a
S1
S3
S5
S8
Stimulus
Rest
Amplitude frequncy (Hz)
Phase frequncy (Hz)
b
S2
S4
S7
S9
S10
S11
S12
Stimulus
Rest
Amplitude frequncy (Hz)
Phase frequncy (Hz)

c

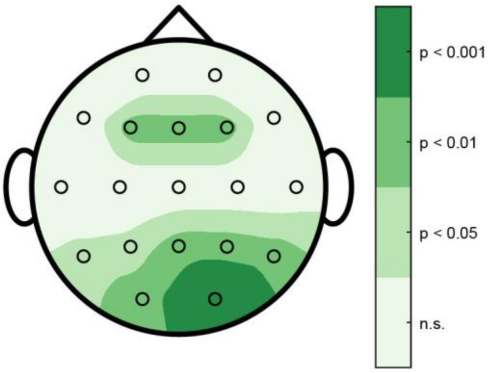


**Supplementary figure 3** Theta-gamma coupling for all participants. (**a**) Comodulograms representing the strength of phase amplitude coupling (PAC) between theta band as the phase frequency and gamma band as the amplitude frequency oscillations based on the mean vector length (MVL) measurement during the stimulus (left) and rest (right) cycles for all participants in the entrained group (n = 4). (**b**) Same as **a** for all participants in the non-entrained group (n = 7). (**c**) Channels with remarkable theta-gamma coupled oscillations on which the MVL difference of the stimulus and rest cycles is significantly different between the two groups. The p-value of the test of significance is color coded. n.s.: not significant.
